# CRISPR-Cas9 enables efficient genome engineering of the strictly lytic, broad host-range staphylococcal bacteriophage K

**DOI:** 10.1101/2024.03.19.585701

**Authors:** Jonas Fernbach, Jasmin Baggenstos, Jeannine Riedo, Shawna McCallin, Martin J. Loessner, Samuel Kilcher

**Author notes:** **Corresponding author**: Samuel Kilcher.

## Abstract

*Staphylococcus aureus* is an important opportunistic pathogen, responsible for a range of diseases that often prove challenging to treat due to resistance to methicillin, vancomycin, and other antimicrobials. Bacteriophages present a promising alternative to target such pathogens, particularly when conventional drugs are ineffective. The antimicrobial efficacy of phage therapeutics can be further improved through genetic engineering. Among *S. aureus* phages, members of the *Twortvirinae* subfamily, characterized by their strictly lytic nature and broad host range, are considered the most promising therapeutic candidates. However, their large genome sizes make them notoriously difficult to engineer. In this study, we utilized *Twortvirus* K as a model to develop an efficient phage engineering platform, leveraging homologous recombination and CRISPR-Cas9-assisted counterselection. As proof of principle, this platform was utilized to construct a nanoluciferase (*nluc*)-encoding reporter phage (K::*nluc*) and tested as a preliminary, bioluminescence-based approach for identifying viable *Staphylococcus* cells. Independent of their phage-resistance profile, 100% of tested clinical *S. aureus* isolates emitted bioluminescence upon K::*nluc* challenge. This diagnostic assay was further adapted to complex matrices such as human whole blood and bovine raw milk, simulating *S. aureus* detection scenarios in bacteremia and bovine mastitis. Beyond reporter phage-based diagnostics, our engineering technology opens avenues for the design and engineering of therapeutic *Twortvirinae* phages to combat drug-resistant *S. aureus* strains.

## Introduction

The prevalence of multi-drug resistant (MDR) pathogens among the human population has been steadily increasing in recent decades [1–4]. The WHO speculates that deaths associated with these MDR organisms might even surpass cancer-related fatalities by 2050 [5]. While small molecule antibiotics have been pivotal in treating widespread diseases, their easy accessibility, over-prescription, and extensive use in medical and agricultural fields have fueled the emergence of resistant strains [1, 6, 7]. Bacteriophages and phage-encoded proteins such as endolysins (enzymes that degrade the bacterial cell wall), present promising alternatives to conventional antibiotic treatment. They are being postulated as a significant force in combating the antibiotic resistance crisis in the coming years [8, 9].

Past case reports demonstrated the successful application of phages to treat MDR infections [10, 11]. Rapid advances in synthetic biology and genetic engineering have allowed for the design, development, and application of genetically altered phage variants with enhanced clinical potential. Furthermore, novel methods for genetically modifying phages to achieve enhanced functionality are constantly being developed. These innovations in phage therapy are evidenced by recent studies and technological advancements which are summarized in [12–15].

Through the strategic selection and engineering of the phage backbone, we can finetune crucial characteristics, such as host specificity, to cater to specific clinical applications. These methods have been employed to transition phages to entirely new hosts [16–19], modify the infection cycle for clinical suitability [20], and deliver antimicrobial payload genes directly to the infection site [21]. The field of synthetic biology-based engineering is also expanding, and it now encompasses a variety of sophisticated DNA manipulation techniques. These include isothermal Gibson assembly [22], yeast-based recombineering [23], and whole-genome synthesis, all of which can be utilized for phage genome modification.

*Staphylococcus aureus*, a prevalent opportunistic pathogen, colonizes up to 50% of humans, and can result in severe diseases such as pneumonia, respiratory-, surgical- and cardiovascular infections, as well as nosocomial bacteremia [24]. Bacteremia alone has an estimated annual incidence of up to 50 cases per 100 000, with 10% to 30% of these patients dying as a direct result of the infection [25]. Furthermore, resistance emergence in *S. aureus* isolates against a number of different antibiotics has been reported. Methicillin-resistant *S. aureus* (MRSA) and vancomycin-resistant *S. aureus* (VRSA) are two of the variants of greatest concern. According to the CDC’s 2019 report, there were more than 323 700 cases of MRSA and 10 600 attributed deaths in 2017 in the United States alone, although the incidence has been decreasing over the years [26].

Bacteriophages from the *Twortvirinae* subfamily, like the model phage K and its close relatives, are ideal candidates for phage therapy. They have a strictly lytic life-cycle and a broad host range across *Staphylococci* [27]. *Staphylococcus* bacteriophage genomes can be intentionally activated for engineering purposes through a couple of methods: transformation of L-form bacteria [28][29] or using non-electroporation *Staphylococcus* transformation (NEST) [30]. These techniques enable the engineering of smaller phage genomes. However, like all synthetic methods, engineering becomes challenging when dealing with larger phage genomes that require assembly from numerous long DNA fragments. We therefore chose a homologous recombination (HR)-based approach, which has been successfully used for engineering larger phage genomes [19, 31–36] and included a downstream CRISPR-Cas-assisted counterselection (CS) step, which allowed us to rapidly obtain recombinant phages. To demonstrate the functionality and applicability of this approach, we engineered a phage K-based reporter phage that enables detection of viable *S. aureus* cells. Reporter phages are engineered to carry a heterologous reporter gene, such as a luciferase, which is expressed during infection. The expression of this gene generates a detectable signal, indicating the presence of viable host cells [37–39]. Reporter phages offer significant advantages for bacterial detection, combining the speed and simplicity of PCR-based methods with the ability to detect viable cells. This contrasts with traditional culture-based methods, which, while reliable for detecting viable cells, are time-consuming and often require several days to yield results [40, 41].

Prior research has demonstrated the successful engineering of strictly lytic, *S. aureus*-infecting reporter bacteriophages through the use of HR [42]. Notably, this work resulted in the positive identification of 97.7% of 390 clinical MRSA isolates at low bacterial concentrations. Recognizing the potential difficulties in the engineering process, our study aimed to devise a strategy that enables the generation of recombinant phages, even those that cannot be isolated through dilution enrichment due to the absence of a detectable reporter protein. This approach is particularly useful for systems characterized by low recombination rates, where traditional methods may prove labor-intensive and time-consuming. To address this, our engineering pipeline incorporates a CRISPR-Cas-based CS system, facilitating the isolation of recombinant phages following HR. The efficacy of this approach has been validated by its successful application in other phage-host systems. [21, 43–48].

Our engineered reporter phage K::*nluc*, was subjected to a comprehensive evaluation involving a diverse panel of 71 *S. aureus* strains, including clinical isolates with varying degrees of vancomycin resistance. Intriguingly, through the use of bioluminescence detection following treatment with K::*nluc*, our approach allowed for the identification of strains that displayed no signs of productive infection when analyzed using conventional infection assays. Furthermore, we demonstrated the functionality of K::*nluc* in complex matrices such as bovine raw milk and human whole blood. These findings underscore the potential of K::*nluc* for the rapid detection of *S. aureus* strains and suggest promising avenues for future research in diagnostic settings.

## Results

### CRISPR-Cas9-assisted engineering of *Staphylococcus* reporter phage K

We established a two-step protocol that includes HR and subsequent CS, as illustrated in **Figure 1A**. In order to circumvent the pervasive restriction barriers often found in many *S. aureus* isolates, we opted to use *S. aureus* RN4220. This laboratory strain, obtained through extensive chemical mutagenesis of strain 8325-4, has had its resident prophages and restriction-modification systems removed. This modification allows for the transformation of RN4220 with *E. coli*-derived plasmid DNA [49]. Phage K, characterized by its broad host range and previously documented therapeutic applications, was selected as our engineering scaffold. Its host range primarily includes *S. aureus* and extends to other *Staphylococcus* species [50], making it a versatile candidate for both diagnostic and therapeutic purposes. This potential is evidenced by successful applications of *Twortvirinae* phages in therapeutic contexts, which have demonstrated their efficacy against a variety of clinically relevant bacterial strains [51–58].

**Figure 1:**
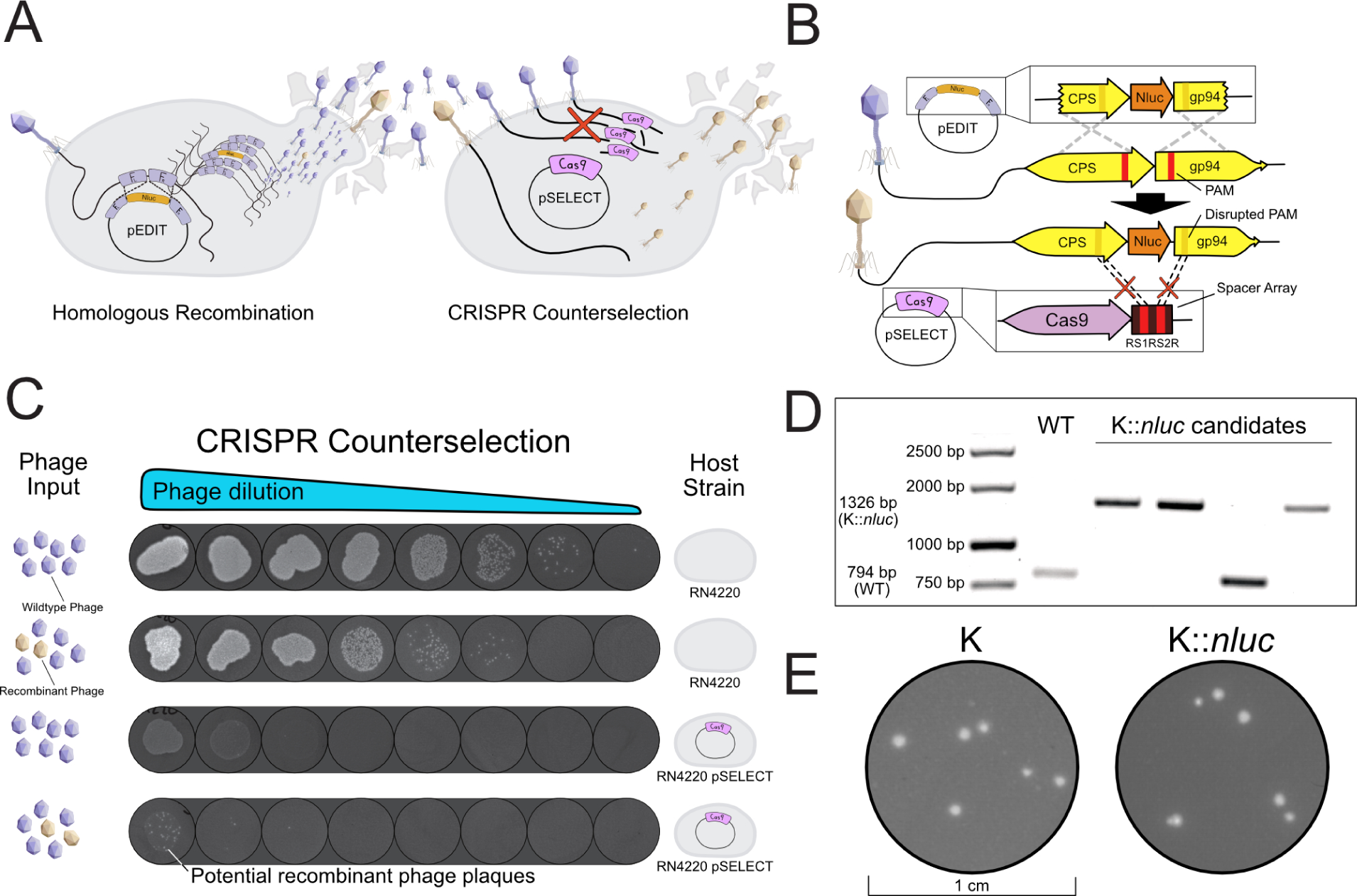
Construction of reporter phage K::*nluc* using a homologous recombination-based and CRISPR-Cas9-assisted phage engineering platform. **(A)** We employ two *S. aureus* bacterial host strains in our process. The first is the recombination donor, RN4220 pEDIT_nluc_ (RN4220 transformed with the HR donor plasmid pEDIT_nluc_), and the second is the counterselection strain, RN4220 pSELECT_CPS_. By sequentially infecting these host strains, we can generate (homologous recombination) and enrich (counterselection) the engineered phage. **(B)** The design of pEDIT_nluc_ and pSELECT_CPS_ facilitates selective amplification of phage particles that have undergone homologous recombination. This is made possible through two synonymous, single-nucleotide polymorphisms in the protospacer adjacent motifs (PAM) of the donor template. The Cas9-mediated restriction of wildtype phage DNA is guided by two spacer sequences on pSELECT_CPS_, flanked by repeat regions (RS1RS2R), which target each homology arm on the target phage genome. **(C)** The efficiency of the CRISPR-Cas9 counterselection system is demonstrated. Wildtype lysate or recombination lysate was serially diluted and spotted on wildtype RN4220 (top two rows) or on the counterselection strain (bottom two rows). While wildtype phages were completely restricted (row three, limit of detection: 100 PFU/ml), visible plaques at the lowest dilutions in row four indicate the presence of phage variants that have escaped CRISPR-Cas9 restriction. **(D)** Individual plaques were selected, and the presence of an insertion of the intended size was validated using PCR. Positive PCR products were then Sanger sequenced to determine the correct genotype at the insertion site. **(E)** Plaque morphologies of K and K::*nluc*.

Utilizing existing data on genome structure and transcriptomic profiles, we identified a region associated with the major capsid protein (*cps*) as a favorable locus for payload insertion. This area has been previously demonstrated to be highly expressed in phage K [59], and it has been a common choice for payload insertion in other phages [21, 31, 42]. To mediate expression from a strong, endogenous promoter, the reporter gene and a ribosomal binding site were integrated immediately downstream of the *cps* coding sequence. We opted for the nanoluciferase (*nluc*) reporter gene, drawing on previous work where it was successfully incorporated into phage K, allowing for the relative quantification of payload production [42].

The first step of the pipeline involves propagating phage K in the presence of the homology donor plasmid (pEDIT_nluc_). This plasmid carries the payload gene, along with up- and downstream flanking homology arms, which guide the sequence-specific integration of the payload. As our payload, we selected a bioluminescent luciferase derived from *Oplophorus gracilirostris*, NanoLuc^®^ (Promega). Despite its relatively small size of merely 516 bp, NanoLuc^®^ produces a detectable signal even at very low concentrations, making it a more sensitive choice compared to other luciferases [60].

The lysate of phage K, produced on RN4220 pEDIT_nluc_, comprises a heterogeneous population, predominantly of wildtype phages, interspersed with a minority of recombinants. This mixed lysate was subsequently propagated on another RN4220 host, one equipped with a counterselection plasmid (pSELECT_CPS_). This plasmid encodes an episomal CRISPR-Cas system, serving as the principal mechanism of selection to isolate the desired recombinant phages.

The pSELECT_CPS_ plasmid encodes the *S. pyogenes cas9* gene, along with tracrRNA and crRNA elements. The crRNA element carries a repeat-spacer1-repeat-spacer2-repeat (RS1RS2R) sequence, with each spacer targeting one homology arm flanking the *nluc* integration site. We designed the protospacer-adjacent-motifs (PAMs) in the editing template (pEDIT_nluc_) to contain silent PAM mutations. This design enables K::*nluc* replication in the presence of pSELECT_CPS_ (**Figure 1B**). Infection of RN4220 pSELECT_CPS_ with wildtype phage K resulted in complete phage restriction (**Figure 1C**). When the same host strain was infected with a lysate containing a mixed population of recombinant and wildtype phages following HR, escape mutants were obtained at a frequency of approximately 10*^−^*^4^ (**Figure 1C**). We then performed PCR amplification of the insertion site, and found that three out of the four plaques we picked yielded PCR products of the expected size (1326bp, **Figure 1D**), indicating successful payload integration. Further confirmation was obtained through Sanger sequencing of the PCR products with the correct band size, which revealed a markerless insertion of the payload at the intended locus.

### Characterization of K::*nluc* infectivity and bioluminescence kinetics across clinically relevant *S. aureus* strains

There was no evident difference in plaque morphology between the wildtype and K::*nluc* phages when infecting RN4220 (**Figure 1E**). We proceeded to assess the infectivity of K::*nluc* across a diverse set of 71 different *S. aureus* strains using efficiency of plaquing (EOP) assays (**Suppl. Figure S1**). This set included the phage propagation host PSK, common laboratory strains, and clinical isolates with varying degrees of vancomycin resistance. Out of the 71 strains tested, 51 showed plaque formation (limit of detection: 100 PFU/ml) when treated with K::*nluc*, indicating successful infection by the phage. This assessment is crucial as it provides insight into the potential range of application for K::*nluc* in detecting different *S. aureus* strains.

Next, we established the minimum inhibitory concentration (MIC) of vancomycin for all strains, utilizing the cutoffs as defined by [61]: vancomycin-susceptible (VSSA) strains exhibited an MIC ≤ 2 µg mL*^−^*^1^, vancomycin intermediate resistant (VISA) strains showed an MIC between 4 µg mL*^−^*^1^ and 8 µg mL*^−^*^1^, and vancomycin resistant (VRSA) strains had an MIC ≥ 16 µg mL*^−^*^1^ (**Suppl. Figure S2**). We selected one phage K-susceptible isolate from each category (PSK (VSSA), LI6 (VISA), VRSA7 (VRSA)), and investigated the kinetics of phage K::*nluc*-induced bioluminescence emission.

In all three K::*nluc* infections (PSK, LI6, VRSA7), we observed similar signal intensities and kinetics of bioluminescence generation. There was a swift and consistent increase until it reached a plateau, with a peak fold-change of approximately 1 × 10^6^ relative light units (RLU) above the background luminescence, achieved roughly after 3 hours (**Figure 2A**). To further assess the sensitivity of our reporter phage system, we examined the minimum dose response of bioluminescence emitted by PSK, LI6, and VRSA7 upon infection with a fixed concentration of K::*nluc* (5×10^7^ PFU*/*mL) (**Figure 2B**). Although the minimum dose response of LI6 (2617 CFU*/*mL) was marginally higher than that of PSK (1093 CFU*/*mL) and VRSA7 (707 CFU*/*mL), all values were within a range comparable to those previously reported for other reporter phages [42, 60, 62].

**Figure 2:**
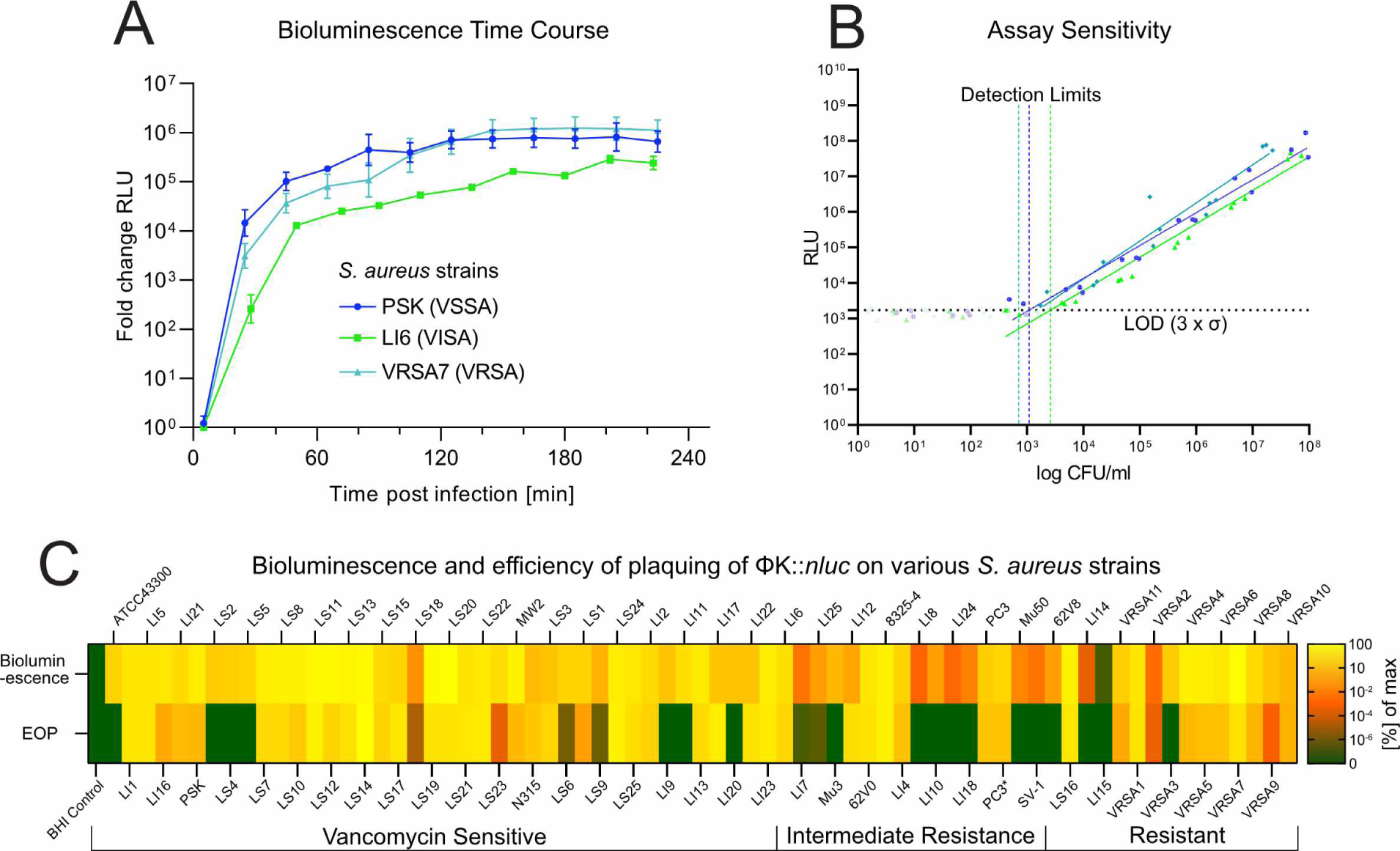
Phage K::*nluc* enables rapid and sensitive detection of a broad range of *S. aureus in vitro*. **(A)** Bioluminescence time course measurements for *S. aureus* strains PSK, LI6 and VRSA7 were obtained by calculating the fold change in relative light units (RLU) compared to infection with K wildtype at a bacterial density of *OD*_600_ = 0.01 and phage titer of 5 × 10^7^ PFU/ml. Values are corrected for background luminescence. Data is mean +/- standard deviation of biological replicates (n=3). **(B)** Minimal dose response of PSK, LI6 and VRSA7 to K::*nluc* was determined by measuring the RLU after 3h of infection for varying bacterial concentrations. The detection limits (vertical dotted lines) were calculated as the minimum cell number required to produce a signal that is higher than three standard deviations (*σ*) above the background luminescence (horizontal dotted lines). Measurements below this cutoff were excluded from linear regression. **(C)** Bioluminescence and efficiency of plaquing (EOP) for 71 *S. aureus* isolates. Bioluminescence for each strain is represented as the mean measured bioluminescence 3h post infection (n=3). EOP is given by the number of plaque forming units (PFU) after infection of a specific strain with K::*nluc*. Values are relative to the host with the highest measurement (LS20 for bioluminescence, LS14 for EOP).

Subsequently, we quantified the range of bioluminescence detection of K::*nluc* across all 71 *S. aureus* strains and compared it with the previously determined EOP (**Figure 2C**). We successfully detected bioluminescence above background levels in all tested strains (71/71), encompassing even those 20 strains that had previously exhibited no evident plaque formation.

### K::*nluc* detection of vancomycin susceptible (VSSA), resistant (VRSA), and intermediate resistant (VISA) *S. aureus* in human whole blood and bovine raw milk

Bacteremia accounts for a significant proportion of bloodstream infections [63, 64]. Considering the complexity of blood as a biological matrix compared to standard laboratory culture conditions, we tested our reporter phage’s functionality in whole human blood. It’s essential to evaluate this, as the *in vitro* infection kinetics of K::*nluc* with pathogenic bacterial strains might differ in more complex biological settings [65–67]. We optimized the assay conditions using whole human blood spiked with the PSK strain, as depicted in **Figure S3**. The most rapid and robust signal response was achieved under the following conditions: the spiked blood was diluted five-fold in growth medium, and the cells were incubated at 37°C for 1 hour to stimulate host cell metabolism before reporter phage infection. We observed higher bioluminescent signals when using a citrate-based anti-coagulant (Na3-citrate, citric acid, glucose, potassium sorbate) compared to Li-Heparin, both of which are common storage solutions in clinical practice.

Leveraging these optimized conditions, we determined the detection limits using the previously selected *S. aureus* strains PSK, LI6, and VRSA7 (**Figure 3A**). These strains were detectable at concentrations as low as 2151, 136, and 1270 CFU/ml, respectively, which is comparable to the results obtained in growth medium (**Figure 2B**).

**Figure 3:**
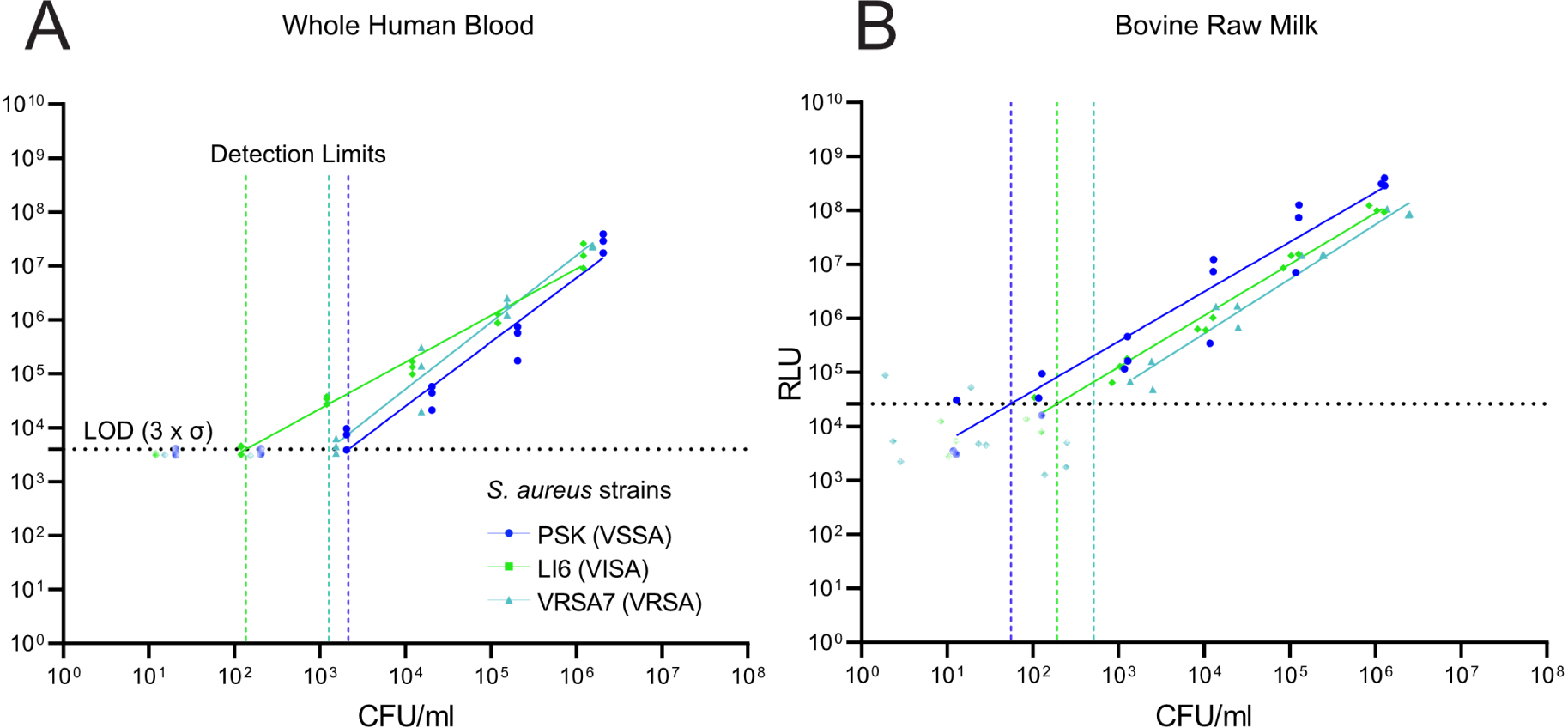
Detection of vancomycin resistant *S. aureus* in whole human blood and bovine raw milk. Minimal dose response of PSK, LI6 and VRSA7 to K::*nluc* in whole human blood **(A)** and bovine raw milk **(B)** was determined by measuring the RLU after 3h of infection for varying bacterial concentrations. Values below the determined limit of detection, set at 3 standard deviations of the mean background luminescence, were excluded. The detection limits (vertical dotted lines) were calculated as the minimum cell number required to produce a reliable signal (three standard deviations (*σ*)) above the mean background luminescence (horizontal dotted line).

Bovine raw milk, another complex matrix, has also been reported to negatively impact bacteriophage proliferation [68–70]. To investigate this, we conducted a similar experiment, replacing blood with unpasteurized, raw bovine milk. Contrary to expectations, treating bovine raw milk infected with *S. aureus* strains PSK, LI6, and VRSA7 with K::*nluc* resulted in comparable levels of bioluminescence and minimal dose responses of 55, 191, and 514 CFU/ml, respectively. This indicates that our optimized setup enables reliable detection of *Staphylococci* by K::*nluc* in human blood and bovine raw milk.

## Discussion

In the face of rising antimicrobial resistance, bacteriophage therapy has garnered increasing attention as a potential alternative to traditional antibiotics [71]. The engineering of bacteriophages has opened up new possibilities, for diagnostic applications and enhancing clinical efficacy [12, 37]. While there are numerous examples of engineered bacteriophages, the engineering of large, strictly lytic, *S. aureus*-infecting bacteriophages has been less explored. A previous study has successfully incorporated *nluc* into two *S. aureus*-infecting phages, ISP and MP115 [42]. ISP and MP115 have been utilized in applications such as phage therapy and Food and Drug Administration (FDA)-approved KeyPath MRSA/methicillin-susceptible *Staphylococcus* aureus (MSSA) assays, respectively [42, 72, 73]. Our study made use of phage K, a member of the Kayvirus subfamily within the *Twortvirinae*, which is recognized for its well-characterized biology, including a transcriptional landscape analysis [59, 74]. We developed a CRISPR-Cas-assisted engineering system for the rapid and reliable generation of genetically altered variants of phage K. As proof of concept, we utilized this pipeline to engineer a phage K mutant containing a bioluminescent *nluc* reporter gene, K::*nluc*.

Given the transformation barriers encountered with clinical *S. aureus* isolates, our current pipeline is constrained to phages capable of infecting restriction-deficient *Staphylococcus* strains. To engineer other phages, a host compatible with the transformation of pEDIT and pSELECT vectors is necessary. For bacteriophages that do not infect RN4220, our approach would necessitate finding a suitable, transformable host, potentially requiring additional steps to remove restriction-modification systems and resident prophages, akin to the modifications made to RN4220 [49, 75]. An alternative approach is the cloning in dcm-*E.coli* strains that carry artificial modification systems [76]. By acknowledging these challenges and implementing effective workarounds, we hope to broaden the applicability of our engineering strategy in the future.

Our engineering workflow was validated through the construction of K::*nluc*, a reporter gene-coding phage K variant for viable *S. aureus* detection. The bioluminescent nanoluciferase, NanoLuc^®^, has been utilized in prior reporter phage development studies [35–37, 59][42, 60, 77–79]. Unlike antimicrobial payloads that disrupt key metabolic pathways and alter bacterium-phage dynamics, intracellular expression of NanoLuc^®^ is less likely to impose significant fitness costs on the phage or host bacterium. Thus, using *nluc* as a payload offered a simple benchmark for our engineering pipeline that was unlikely to affect the fitness of phage K. Given its large genome size (148 kb) and terminal redundancy, phage K tolerates the integration of the relatively small (516 bp) *nluc* payload without inducing genome packaging defects. We postulate that the integration of even larger payloads could be feasible, expanding the scope of potential applications, although a systematic analysis of such scenarios was not conducted in our study.

In standard spot-on-lawn infection screens, 20 out of the 71 bacterial strains did not exhibit plaque formation of K::*nluc*, yet bioluminescence was detected in all strains. This observation suggests that phage binding, DNA delivery, and gene expression are still occurring, even in the absence of visible plaque formation. In such scenarios, it is likely that the infection cycle is interrupted—possibly by the cell undergoing abortive infection—before the completion of lysis and the release of progeny phage particles. For diagnostic applications, this implies that partially phage-resistant strains can be detected successfully as well, albeit with somewhat reduced sensitivity. At the same time, our data demonstrates that a single Kayvirus scaffold could be used to deliver therapeutic effector genes to most *S. aureus* isolates.

The composition of blood can influence phage infection kinetics. At the same time, bacteriophages may be inactivated by innate or adaptive immune responses. [65–67]. Given these factors, it is imperative to acknowledge that the *in vitro* infection kinetics observed may not directly correlate to more complex biological environments. Thus, each reporter phage assay needs to be tailored to the specific matrix or application where a diagnostic need is identified. Within the scope of *S. aureus*, our focus is on bacteremia and bovine mastitis, representing pertinent diseases in humans and animals, respectively. For phage K, it has been previously demonstrated that the presence of Whey proteins in bovine raw milk can competitively inhibit the phage’s attachment to the cell surface, consequently significantly diminishing infection efficiency [68, 70]. Notably, in our experiments, there was no difference in observed detection limits. This may be attributed to matrix dilution, to the added 1 hour activation step, or potentially to a low baseline concentration of whey proteins present in our milk sample.

While the potential of engineered bacteriophages in the treatment and diagnostics of *S. aureus* infections is well-acknowledged, their immediate implementation is hampered by several challenges. For instance, the swift progression of severe symptoms in cases of bacteremia, leading to sepsis, often mandates immediate intervention with broad-spectrum antibiotics, without preliminary identification of the causative pathogen. Nonetheless, the growing inclination towards patient-specific treatments and precision medicine is paving the way for the application of engineered phages. Specifically, our pipeline could facilitate the engineering of *S. aureus*phages to express antimicrobial effector genes, presenting a promising avenue for the future treatment of chronic infections such as wounds, pulmonary conditions, and implant infections, or as a last resort for bacteremia treatment when antibiotics are ineffective.

Additionally, the broad infectivity of K::*nluc* provides significant advantages by enabling the detection of a wide range of strains. However, it also brings forth challenges that need thoughtful consideration, particularly the risk of false positive detections of other *Staphylococcus* species. *Staphylococcus epidermidis*, commonly found in human skin microbiota [80], serves as a prime example. Several strains of *S. epidermidis* have shown susceptibility to phage K infection, underscoring the importance of careful interpretation of detection results across diverse biological samples [50].

Finally, while the interaction of bacteriophages with dormant bacterial cells is complex and not fully understood, the potential for reporter phages to interact with such cells presents an intriguing area for future investigation. For *E. coli* and *P. aerigunosa*, there are instances where phages have been observed to adsorb to dormant hosts and deliver their genome, albeit subsequently entering a state of hibernation or pseudolysogeny, respectively. [81, 82]. Recent studies have also reported the phage infection and lysis of *P. aeruginosa* dormant persister cells [83]. Therefore, future studies could explore the potential of K::*nluc* to interact with and potentially detect dormant *S. aureus* cells, contributing to a more comprehensive understanding of bacteriophage-host dynamics.

In summary, our study presents a novel and validated engineering pipeline for the generation of genetically altered phage K variants, offering promising avenues for *S. aureus* detection and treatment. While challenges remain in translating these findings to complex biological settings, the insights gained here lay the groundwork for further exploration and optimization, potentially contributing to the growing arsenal of tools in the fight against antimicrobial resistance.

## Materials and Methods

### Vancomycin minimum inhibitory concentration (MIC) of selected *S. aureus* strains

The minimum inhibitory concentration (MIC) of the 71 *S. aureus* strains used in this study was determined under the standard culture conditions as outlined in [61]. In brief, 96 well plates were prepared, each well containing 250 µL of Miller-Hinton broth supplemented with a range of vancomycin concentrations (from 0.0625 µg/mL to 64 µg/mL, with a twofold increase between subsequent concentrations). Each well was inoculated with 2 µL of 1:1000 diluted bacterial cultures (grown overnight at 37 *^◦^*C) for each of the 71 strains. After an incubation period of 18 h at 37 *^◦^*C, turbidity was measured photometrically (*OD*_600_). Wells with an *OD*_600_ > 0.1, were considered turbid. The strains were then classified as vancomycin-susceptible (VSSA, MIC ≤ 2 µg/mL), vancomycin intermediate resistant (VISA, MIC = 4 µg/mL to 8 µg/mL), or vancomycin resistant (VRSA, MIC ≥ 16 µg/mL) based on the MIC values.

### Bacterial strains and culture conditions

*S. aureus* PSK (ATCC 19685) was used as propagation and *S. aureus* RN4220 (DSM 26309) as the engineering host of K and K::*nluc*. *E. coli* XL1-blue MRF’ (Stratagene) was used for plasmid amplification prior to RN4220 transformation. RN4220 and XL1-blue cultures were grown overnight (O/N) at 37 *^◦^*C in Brain Heart Infusion (BHI) Broth (Biolife Italiana) and Luria-Bertani/Lysogeny broth (LB) medium (3 M sodium chloride, 10 g/L tryptone, 5 g/L yeast extract, pH 7.2), respectively. The selection of 71 laboratory strains and clinical isolates (Figure S2) were grown on BHI Broth with corresponding antibiotic supplements.

### Bacteriophage propagation

Phage K was propagated on *S. aureus* PSK using the soft-agar overlay method with BHI as bottom agar and LC agar (LB supplemented with 10 mM CaCl_2_, 10 mM MgSO_4_, 10 g/L glucose) as top agar. Overlays were incubated O/N at 37 *^◦^*C and phages extracted from plates with semi-confluent lysis using 5 mL SM buffer (4 *^◦^*C, 2 h, constant agitation). Lysates were sterile-filtered (0.22 µm pores). Phage particles were precipitated O/N at 4 *^◦^*C using polyethylene-glycol (7 % PEG8000 and 1 M NaCl) and purified using stepped CsCl gradient ultracentrifugation. The obtained phage suspension was dialyzed twice against 1000x excess of SM buffer. Purified samples were then stored long-term at 4 *^◦^*C. The titer was determined using the soft-agar overlay method.

### Electroporation of bacterial strains

XL1-blue electrocompetent cells were electroporated at 2.5 kV, 200 Ω, 25 µF, incubated for 1 h with SOC recovery medium (2 % (w/v) Tryptone, 0.5 % (w/v) Yeast extract, 10 mM NaCl, 2.5 mM KCl, 10 mM MgCl_2_, 20 mM Glucose) at 37 *^◦^*C, and plated on selective agar to isolate successful transformants. RN4220 electrocompetent cells were electroporated at 1.8 kV, 600 Ω, 10 µF with 500 ng to 1000 ng of amplified plasmid DNA, incubated for 1 h at 37 *^◦^*C in B2 recovery medium (10 g/L casein hydrolysate, 5 g/L D-glucose, 1 g/L potassium phosphate dibasic, 25 g/L NaCl, 25 g/L yeast extract) and plated on selective medium to isolate successful transformants.

### Plasmid design

The plasmid pLEB579 (kindly gifted by T. Takala, University of Helsinki, Finland) is a shuttle vector shown to have high transformation efficiencies in both *E. coli* and *S. aureus* and was therefore used as a backbone for both the editing template (pEDIT_nluc_) and the CRISPR-Cas9-counterselection system (pSELECT_CPS_). We used a previously reported, *Streptococcus pyogenes*-derived Cas9 (SpyCas9)-based CRISPR system [21] and exchanged the two spacers in the repeat-spacer1-repeat-spacer2-repeat (RS1RS2R) region. This was done to allow targeted restriction of wildtype phage K at two distinct loci (8304 bp and 148 bp up- and downstream of the intended insertion site, respectively) designed to contain a PAM-disrupting synonymous mutation in the successful recombinants. pEDIT_nluc_ was constructed by integrating an nluc gene (optimized for *S. aureus* codon usage, avoidance of Rho-independent termination and an added upstream ribosome binding site: GAGGAGGTAAATATAT), flanked by 400 bp (upstream) and 300 bp (downstream) homology arms, corresponding to the intended K insertion site, into the linearized pLEB579 backbone. Two silent point mutations were included to disrupt the two PAMs adjacent to the target DNA (**Figure 1B**). All synthetic sequences were acquired as GeneArt String DNA Fragments (Thermo Fischer), albeit the RS1RS2R sequences, which were ordered as GeneArt Gene Synthesis (Thermo Fischer) (**Table S2**). Assembly of all constructs was performed using isothermal Gibson Assembly^®^ [22] (NEBuilder^®^ HiFi DNA Assembly Master Mix) and subsequently transformed into *E. coli* XL1-blue MRF’ for plasmid amplification. Plasmids and primers are listed in (**Table S1**).

### CRISPR-Cas9-assisted engineering of K::*nluc*

pEDIT_nluc_ and pSELECT_CPS_ were transformed into *S. aureus* RN4220 to acquire the strains required for recombination and counterselection, respectively. RN4220 pEDIT_nluc_was infected with K via soft-agar overlay and a high titer lysate obtained as described previously (see bacteriophage propagation). Plates showing semi-confluent lysis were washed with SM buffer and 10-fold dilutions of the resulting lysate were used to perform soft-agar overlays on RN4220 pSELECT_CPS_. Individual plaques were picked from the plates showing the fewest (non-zero) plaques, resuspended in SM buffer, and clonally isolated by three rounds of plaque-purification. PCR amplification using primers flanking the insert site (Table S1) was performed and products showing a size indicative of the intended insertion purified and Sanger sequenced (Microsynth AG, Balgach, Switzerland) to validate the correct genomic sequence. Validated phage lysates were purified using ultracentrifugation on a caesium-chloride gradient and subsequent dialysis as described in the section bacteriophage propagation.

### Soft-agar overlay

5 ml BHI soft-agar were melted and cooled to 47 *^◦^*C. The molten soft-agar was inoculated with 200 µL bacterial culture of adequate turbidity (*OD*_600_ > 1) and 10 µL of phage suspension, briefly mixed by agitation, and spread evenly onto BHI agar plates with. Plates were let dry for 15 minutes at room temperature (RT) and subsequently inverted and incubated at 37 *^◦^*C for 12 h to 18 h.

### Spot-on-lawn assay

5 ml BHI soft-agar were melted and let cooled to 47 *^◦^*C. The molten soft-agar was inoculated with 200 µL of bacterial culture, briefly mixed by vortexing, and spread evenly on BHI agar plates. Plates were dried at RT for 15 minutes. 10 µL droplets of phage suspension were then placed carefully on the dried soft-agar. Plates were dried for 15 min, inverted, and incubated at 37 *^◦^*C for 16 h.

### Determination of efficiency of plaquing (EOP)

EOP of bacteriophage suspensions was determined by performing spot-on-lawn assays on bacterial strains of interest. 10-fold dilutions of the phage suspension were prepared up to a maximum dilution of 1 × 10*^−^*^8^. Spot-on-lawn assays were performed as described in the previous section using the series of bacteriophage dilutions and O/N cultures of the corresponding host strains. The EOP is given by the number of plaque forming units (PFU) after infection of a specific strain with K::*nluc*. Values are relative to the host with the highest measurement (LS20 for bioluminescence, LS14 for EOP).

### Bioluminescence time course assay

Stationary phase bacterial cultures were diluted to *OD*_600_ = 0.01, inoculated with 5 × 10^7^ PFU/mL K::*nluc* and incubated at 37 *^◦^*C (180 rpm agitation). Bioluminescence measurements were taken by combining 25 µL of the sample solution with an equal volume of prepared buffer-reconstituted nluc substrate as detailed by the manufacturer (NanoGlo Luciferase Assay System; Promega). Measurements were taken every 20 min (225 min total) in Nunc™ F96 MicroWell™ 446 plates using a GloMax^®^ navigator luminometer (Promega) with 5 s integration time and 2 s delay. To determine the background-corrected fold-change in relative light units (RLU), measurements were normalized to a control reaction of K::*nluc* in BHI medium and the fold-change was calculated as the difference in RLU to an infection of the same strain with wildtype K.

### Minimal dose-response

To determine the minimum concentration of cells giving a significant bioluminescence signal above the background, stationary phase bacterial cultures were diluted to a range of concentrations in ten-fold increments (*OD*_600_ = 10*^−^*^1^ to 10*^−^*^10^). Measurements were taken 3 h post-infection (5 × 10^7^ PFU/mL) and background corrected as described in the previous section. The minimum concentration of cells that could be detected was established as the smallest measurement exceeding a signal threshold, defined as the mean plus three times the standard deviation of the background signal. The theoretical minimal concentration was ascertained at the point where the established signal threshold intersects with a linear regression of the data, considering only RLU values surpassing the threshold.

### Determination of bioluminescence for the 71 *S. aureus* strains used in the study

Stationary phase bacterial cultures were diluted to *OD*_600_ = 0.01, inoculated with 5 × 10^7^ PFU/mL K::*nluc* and incubated at 37 *^◦^*C (180 rpm agitation). Fold-change bioluminescence was measured at 3 hours post-infection and background corrected.

### K::nluc-based detection of *S. aureus* in patient blood and bovine raw milk

Minimal dose response of K::*nluc* infection on one representative each of VSSA, VISA and VRSA was conducted in triplicate as done with regular growth medium described above, albeit with some modifications. First, spiked whole human blood or bovine raw milk were mixed 1:5 with BHI growth medium and incubated 1 h at 37 *^◦^*C with agitation (180 rpm) prior to infection with 5×10^7^ PFU/mL K::*nluc*. Bioluminescence was measured after 3 h. The whole human blood samples were stored in anticoagulant solutions containing either 1.89 mg/mL Na3-citrate, 0.69 mg/mL citric acid, 2.1 mg/mL glucose and 0.03 mg/mL potassium sorbate (BD Vacutainer^®^ (REF 367756), Becton, Dickinson and Company, New Jersey, USA) or Li-Heparin (17 IU/mL) (BD Vacutainer^®^ (REF 368886), Becton, Dickinson and Company, New Jersey, USA).

### Software

CLC Genomics Workbench version 20.0.4 was used for sequence analyses such as primer and string design as well as evaluation of sanger sequencing results. Plotting was done using GraphPad Prism version 10.0.0 for Windows. OpenAI’s ChatGPT 4 [84] was used as a tool to assist with formatting and editing the manuscript. This involved iterative refinements to ensure clarity and conciseness of the content presented.

## Supplementary Material

**Supplementary Figure 1:**
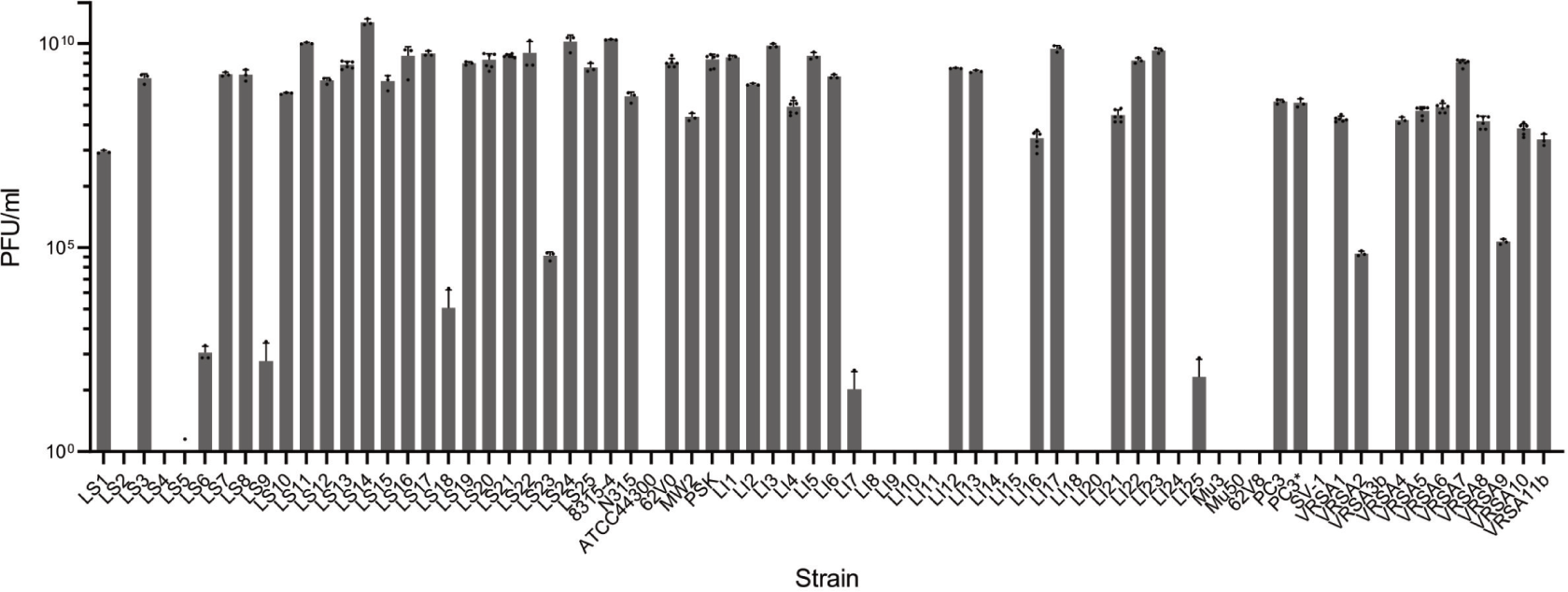
Efficiency of plating (EOP) of K::*nluc* on a selection of 71 *S. aureus* species. Standard spot-on-lawn plaque assay was performed to elucidate the absolute EOP of K::*nluc* on a panel of 71 *S. aureus* species given by plaque forming units per volume (PFU/ml). Standard deviation was calculated from 3-6 replicates per strain.

**Supplementary Figure 2:**
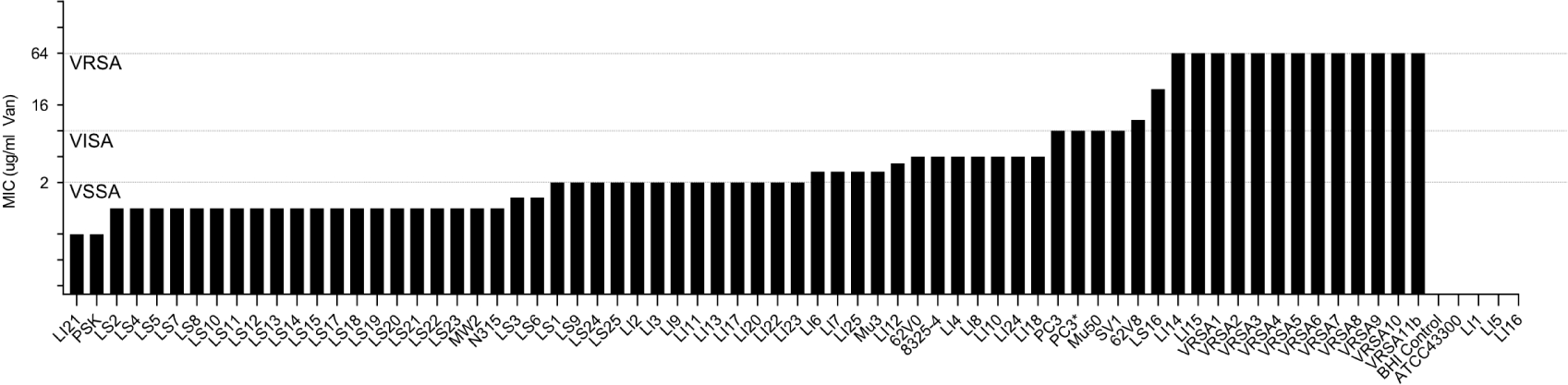
Minimal inhibitory concentration (MIC) of vancomycin on bacterial strains used in this study. The minimum inhibitory concentration (MIC) of vancomycin on the 71 bacterial strain used in this study was tested by culturing in the presence of varying vancomycin concentrations, ranging from 0.0625 µg/mL – 64 µg/mL with twofold increase between subsequent concentrations. For cases where replicates did not yield uniform results, the MIC was determined as the triplicate mean. Vancomycin suceptibility (VSSA), intermediate resistance (VISA) and full resistance (VRSA) are indicated along with the MIC cutoffs defined by [61].

**Supplementary Figure 3:**
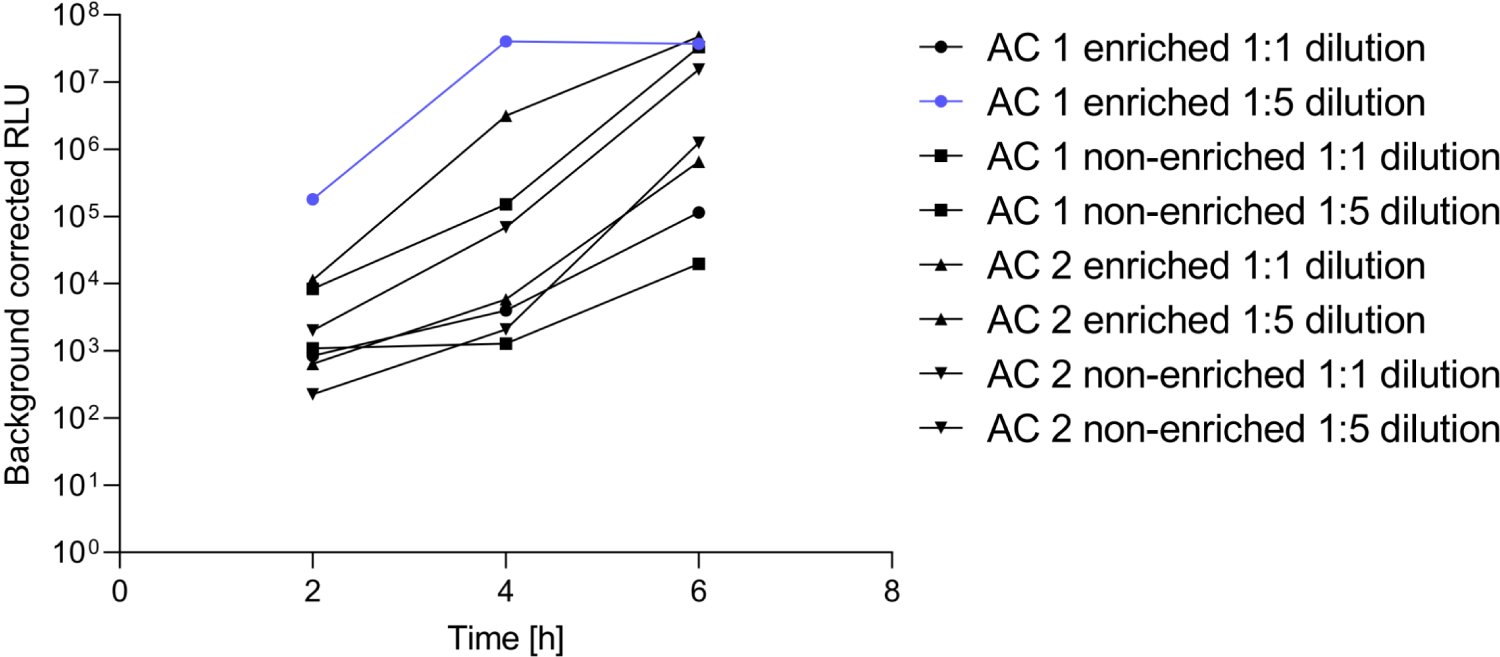
Effects of various parameters on K::*nluc* infection-associated bioluminescence in human whole blood. Whole human blood was spiked with different bacterial concentrations of *S. aureus* PSK. Samples were subsequently diluted with BHI growth medium at a ratio of 1:1 or 1:5. Samples were then either infected with 5 × 10^7^ PFU/ml K::*nluc* or first incubated at 37 *^◦^*C for 1 h prior to infection with K::*nluc*. These parameters were furthermore tested on two types of anticoagulant solutions, AC1 (Na3-citrate, citric acid, glucose, potassium sorbate) and AC2 (Li-Heparin). The combination of parameters resulting in the highest bioluminescence is highlighted in blue.

**Supplementary Table 1:**
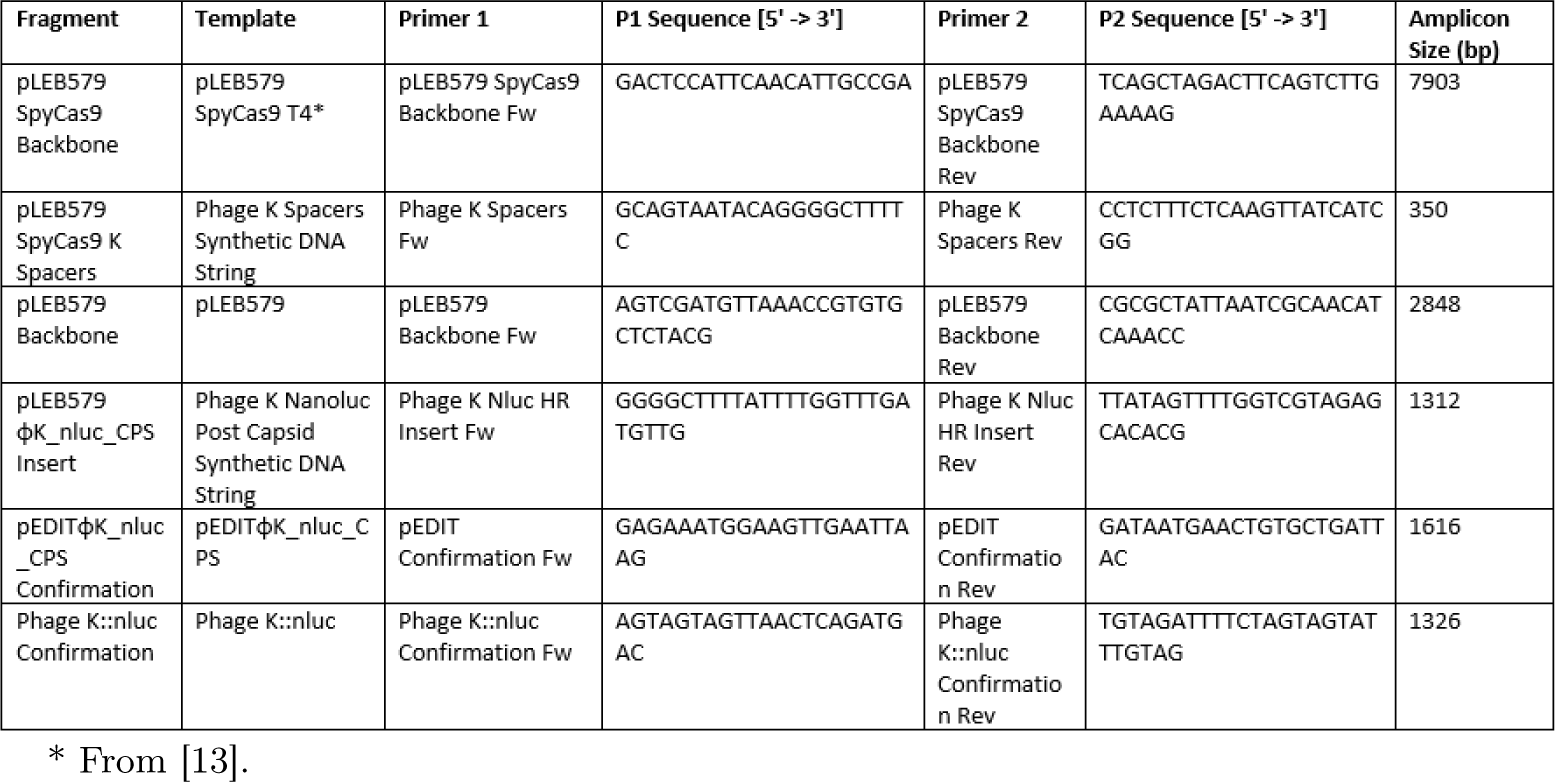
Primers and templates used for plasmid construction and insert confirmations.

**Supplementary Table 2:**
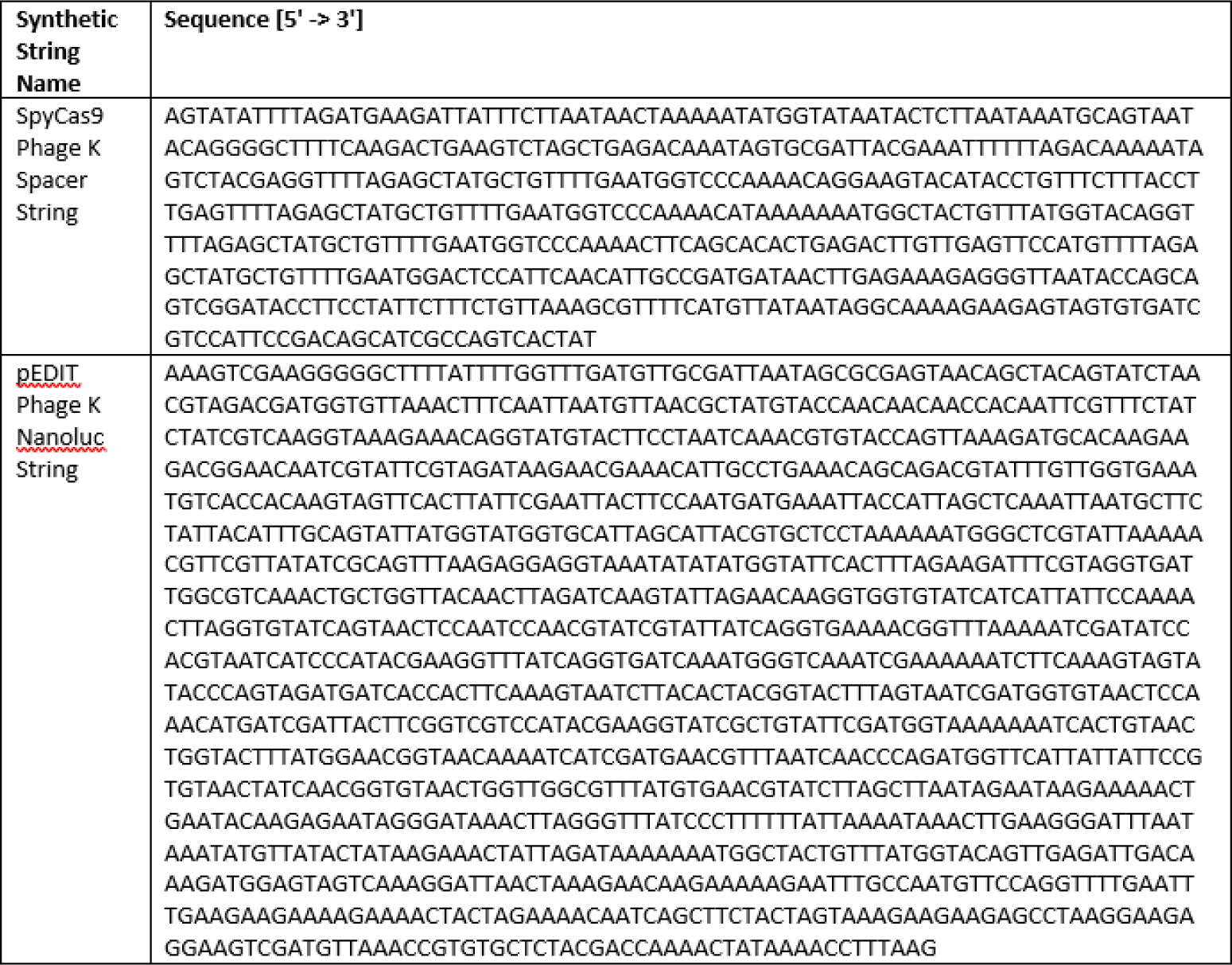
SpyCas9 Phage K spacer string and pEDIT Phage K::*nluc* string used in this study.

## References

1. Aslam, B. et al. Antibiotic resistance: a rundown of a global crisis. Infection and drug resistance, 1645–1658 (2018).

2. De Oliveira, D. M. et al. Antimicrobial resistance in ESKAPE pathogens. Clinical microbiology reviews 33, 10–1128 (2020).

3. Prestinaci, F., Pezzotti, P. & Pantosti, A. Antimicrobial resistance: a global multi-faceted phenomenon. Pathogens and global health 109, 309–318 (2015).

4. Aslam, B. et al. Antibiotic resistance: one health one world outlook. Frontiers in cellular and infection microbiology, 1153 (2021).

5. O’neill, J. Antimicrobial resistance: tackling a crisis for the health and wealth of nations. Rev. Antimicrob. Resist. (2014).

6. Perry, J., Waglechner, N. & Wright, G. The prehistory of antibiotic resistance. Cold Spring Harbor perspectives in medicine 6 (2016).

7. Skandalis, N. et al. Environmental spread of antibiotic resistance. Antibiotics 10, 640 (2021).

8. Hatfull, G. F., Dedrick, R. M. & Schooley, R. T. Phage therapy for antibiotic-resistant bacterial infections. Annual Review of Medicine 73, 197–211 (2022).

9. Anyaegbunam, N. J. et al. The resurgence of phage-based therapy in the era of increasing antibiotic resistance: From research progress to challenges and prospects. Microbiological research, 127155 (2022).

10. McCallin, S., Sacher, J. C., Zheng, J. & Chan, B. K. Current state of compassionate phage therapy. Viruses 11, 343 (2019).

11. Hitchcock, N. M. et al. Current Clinical Landscape and Global Potential of Bacteriophage Therapy. Viruses 15, 1020 (2023).

12. Lenneman, B. R., Fernbach, J., Loessner, M. J., Lu, T. K. & Kilcher, S. Enhancing phage therapy through synthetic biology and genome engineering. Current Opinion in Biotechnology 68, 151–159 (2021).

13. Meile, S., Du, J., Dunne, M., Kilcher, S. & Loessner, M. J. Engineering therapeutic phages for enhanced antibacterial efficacy. Current opinion in virology 52, 182–191 (2022).

14. Mahler, M., Costa, A. R., van Beljouw, S. P., Fineran, P. C. & Brouns, S. J. Approaches for bacteriophage genome engineering. Trends in Biotechnology (2022).

15. Usman, S. S., Uba, A. I. & Christina, E. Bacteriophage genome engineering for phage therapy to combat bacterial antimicrobial resistance as an alternative to antibiotics. Molecular Biology Reports, 1–13 (2023).

16. Tétart, F., Repoila, F., Monod, C. & Krisch, H. Bacteriophage T4 host range is expanded by duplications of a small domain of the tail fiber adhesin. Journal of molecular biology 258, 726–731 (1996).

17. Dunne, M. et al. Reprogramming bacteriophage host range through structure-guided design of chimeric receptor binding proteins. Cell reports 29, 1336–1350 (2019).

18. Yehl, K. et al. Engineering phage host-range and suppressing bacterial resistance through phage tail fiber mutagenesis. Cell 179, 459–469 (2019).

19. Zhang, J. et al. Expansion of the plaquing host range and improvement of the absorption rate of a T5-like Salmonella phage by altering the long tail fibers. Applied and Environmental Microbiology 88, e00895–22 (2022).

20. Dedrick, R. M. et al. Engineered bacteriophages for treatment of a patient with a disseminated drug-resistant Mycobacterium abscessus. Nature medicine 25, 730–733 (2019).

21. Du, J. et al. Enhancing bacteriophage therapeutics through in situ production and release of heterologous antimicrobial effectors. Nature Communications 14, 4337 (2023).

22. Gibson, D. G. et al. Enzymatic assembly of DNA molecules up to several hundred kilobases. Nature methods 6, 343–345 (2009).

23. Jaschke, P. R., Lieberman, E. K., Rodriguez, J., Sierra, A. & Endy, D. A fully decompressed synthetic bacteriophage øX174 genome assembled and archived in yeast. Virology 434, 278–284 (2012).

24. Cheung, G. Y., Bae, J. S. & Otto, M. Pathogenicity and virulence of Staphylococcus aureus. Virulence 12, 547–569 (2021).

25. Van Hal, S. J. et al. Predictors of mortality in Staphylococcus aureus bacteremia. Clinical microbiology reviews 25, 362–386 (2012).

26. Centers for Disease Control and Prevention 2021 Nov. 2021. https://www.cdc.gov/drugresistance/biggest-threats.html.

27. Vandersteegen, K. et al. Romulus and Remus, two phage isolates representing a distinct clade within the Twortlikevirus genus, display suitable properties for phage therapy applications. Journal of virology 87, 3237–3247 (2013).

28. Kilcher, S., Studer, P., Muessner, C., Klumpp, J. & Loessner, M. J. Cross-genus rebooting of custom-made, synthetic bacteriophage genomes in L-form bacteria. Proceedings of the National Academy of Sciences 115, 567–572 (2018).

29. Fernbach, J., Meile, S., Kilcher, S. & Loessner, M. J. in Bacteriophage Therapy: From Lab to Clinical Practice 247–259 (Springer, 2023).

30. Assad-Garcia, N. et al. Cross-genus “boot-up” of synthetic bacteriophage in staphylococcus aureus by using a new and efficient DNA Transformation method. Applied and Environmental Microbiology 88, e01486–21 (2022).

31. Loessner, M. J., Rees, C., Stewart, G. & Scherer, S. Construction of luciferase reporter bacteriophage A511:: luxAB for rapid and sensitive detection of viable Listeria cells. Applied and environmental microbiology 62, 1133–1140 (1996).

32. Namura, M., Hijikata, T., Miyanaga, K. & Tanji, Y. Detection of Escherichia coli with fluorescent labeled phages that have a broad host range to E. coli in sewage water. Biotechnology progress 24, 481–486 (2008).

33. Mahichi, F., Synnott, A. J., Yamamichi, K., Osada, T. & Tanji, Y. Site-specific recombination of T2 phage using IP008 long tail fiber genes provides a targeted method for expanding host range while retaining lytic activity. FEMS microbiology letters 295, 211–217 (2009).

34. Erickson, S. et al. Isolation and engineering of a Listeria grayi bacteriophage. Scientific Reports 11, 18947 (2021).

35. Shitrit, D. et al. Genetic engineering of marine cyanophages reveals integration but not lysogeny in T7-like cyanophages. The ISME journal 16, 488–499 (2022).

36. Guan, J. et al. Bacteriophage genome engineering with CRISPR–Cas13a. Nature microbiology 7, 1956–1966 (2022).

37. Meile, S., Kilcher, S., Loessner, M. J. & Dunne, M. Reporter phage-based detection of bacterial pathogens: design guidelines and recent developments. Viruses 12, 944 (2020).

38. Hussain, W., Ullah, M. W., Farooq, U., Aziz, A. & Wang, S. Bacteriophage-based advanced bacterial detection: Concept, mechanisms, and applications. Biosensors and Bioelectronics 177, 112973 (2021).

39. Ye, J. et al. Phage-based technologies for highly sensitive luminescent detection of foodborne pathogens and microbial toxins: A review. Comprehensive Reviews in Food Science and Food Safety 21, 1843–1867 (2022).

40. Peri, A. M., Harris, P. N. & Paterson, D. L. Culture-independent detection systems for bloodstream infection. Clinical Microbiology and Infection 28, 195–201 (2022).

41. Foddai, A. C. & Grant, I. R. Methods for detection of viable foodborne pathogens: Current state-of-art and future prospects. Applied Microbiology and Biotechnology 104, 4281–4288 (2020).

42. Brown, M. et al. Development and evaluation of a sensitive bacteriophage-based MRSA diagnostic screen. Viruses 12, 631 (2020).

43. Kiro, R., Shitrit, D. & Qimron, U. Efficient engineering of a bacteriophage genome using the type IE CRISPR-Cas system. RNA biology 11, 42–44 (2014).

44. Tao, P., Wu, X., Tang, W.-C., Zhu, J. & Rao, V. Engineering of bacteriophage T4 genome using CRISPR-Cas9. ACS synthetic biology 6, 1952–1961 (2017).

45. Manor, M. & Qimron, U. Selection of genetically modified bacteriophages using the CRISPR-Cas system. Bio-protocol 7, e2431–e2431 (2017).

46. Møller-Olsen, C., Ho, S., Shukla, R., Feher, T. & Sagona, A. Engineered K1F bacteriophages kill intracellular Escherichia coli K1 in human epithelial cells. Sci Rep 8: 17559 (2018).

47. Hupfeld, M. et al. A functional type II-A CRISPR–Cas system from Listeria enables efficient genome editing of large non-integrating bacteriophage. Nucleic acids research 46, 6920–6933 (2018).

48. Corts, A. D., Thomason, L. C., Gill, R. T. & Gralnick, J. A. Efficient and precise genome editing in Shewanella with recombineering and CRISPR/Cas9-mediated counter-selection. ACS Synthetic Biology 8, 1877–1889 (2019).

49. Kreiswirth, B. N. et al. The toxic shock syndrome exotoxin structural gene is not detectably transmitted by a prophage. Nature 305, 709–712 (1983).

50. Göller, P. C. et al. Multi-species host range of staphylococcal phages isolated from wastewater. Nature Communications 12, 6965 (2021).

51. O’flaherty, S. et al. Potential of the polyvalent anti-Staphylococcus bacteriophage K for control of antibiotic-resistant staphylococci from hospitals. Applied and environmental microbiology 71, 1836–1842 (2005).

52. Merabishvili, M. et al. Quality-controlled small-scale production of a well-defined bacteriophage cocktail for use in human clinical trials. PloS one 4, e4944 (2009).

53. Lehman, S. M. et al. Design and preclinical development of a phage product for the treatment of antibiotic-resistant Staphylococcus aureus infections. Viruses 11, 88 (2019).

54. Łubowska, N. et al. Characterization of the three new kayviruses and their lytic activity against multidrug-resistant Staphylococcus aureus. Microorganisms 7, 471 (2019).

55. Van Nieuwenhuyse, B. et al. A case of in situ phage therapy against Staphylococcus aureus in a bone allograft polymicrobial biofilm infection: Outcomes and phage-antibiotic interactions. Viruses 13, 1898 (2021).

56. Kolenda, C. et al. Development of Phage Therapy to Treat Staphylococci Bone and Joint Infections in France: Isolation and Characterization of 17 Novel Anti-Staphylococcus Bacteriophages. 103, 86–86 (2021).

57. Onsea, J. et al. Bacteriophage therapy for the prevention and treatment of fracture-related infection caused by Staphylococcus aureus: a preclinical study. Microbiology spectrum 9, e01736–21 (2021).

58. Plumet, L. et al. Bacteriophage therapy for Staphylococcus aureus Infections: A review of animal models, treatments, and clinical trials. Frontiers in Cellular and Infection Microbiology, 808 (2022).

59. Finstrlová, A. et al. Global transcriptomic analysis of bacteriophage-host interactions between a Kayvirus therapeutic phage and Staphylococcus aureus. Microbiology spectrum 10, e00123–22 (2022).

60. Meile, S. et al. Engineered reporter phages for rapid bioluminescence-based detection and differentiation of viable Listeria cells. Applied and environmental microbiology 86, e00442–20 (2020).

61. Clinical & Institute, L. S. Performance Standards for Antimicrobial Susceptibility Testing 28th ed. (Clinical and Laboratory Standards Institute, Wayne, PA, 2018).

62. Kim, S., Kim, M. & Ryu, S. Development of an engineered bioluminescent reporter phage for the sensitive detection of viable Salmonella typhimurium. Analytical chemistry 86, 5858–5864 (2014).

63. Reimer, L. G., Wilson, M. L. & Weinstein, M. P. Update on detection of bacteremia and fungemia. Clinical microbiology reviews 10, 444–465 (1997).

64. Viscoli, C. Bloodstream infections: the peak of the iceberg 2016.

65. Srivastava, A. S., Kaido, T. & Carrier, E. Immunological factors that affect the in vivo fate of T7 phage in the mouse. Journal of virological methods 115, 99–104 (2004).

66. Principi, N., Silvestri, E. & Esposito, S. Advantages and limitations of bacteriophages for the treatment of bacterial infections. Frontiers in pharmacology 10, 513 (2019).

67. Porayath, C. et al. Characterization of the bacteriophages binding to human matrix molecules. International Journal of Biological Macromolecules 110, 608–615 (2018).

68. O’flaherty, S., Coffey, A., Meaney, W., Fitzgerald, G. & Ross, R. Inhibition of bacteriophage K proliferation on Staphylococcus aureus in raw bovine milk. Letters in applied microbiology 41, 274–279 (2005).

69. Gill, J. et al. Efficacy and pharmacokinetics of bacteriophage therapy in treatment of subclinical Staphylococcus aureus mastitis in lactating dairy cattle. Antimicrobial agents and chemotherapy 50, 2912–2918 (2006).

70. Gill, J., Sabour, P., Leslie, K. & Griffiths, M. Bovine whey proteins inhibit the interaction of Staphylococcus aureus and bacteriophage K. Journal of applied microbiology 101, 377–386 (2006).

71. Abedon, S. T., Garcıa, P., Mullany, P. & Aminov, R. Phage therapy: past, present and future 2017.

72. Vandersteegen, K. et al. Microbiological and molecular assessment of bacteriophage ISP for the control of Staphylococcus aureus. PLoS One 6, e24418 (2011).

73. Bhowmick, T. et al. Controlled multicenter evaluation of a bacteriophage-based method for rapid detection of Staphylococcus aureus in positive blood cultures. Journal of clinical microbiology 51, 1226–1230 (2013).

74. Barylski, J., Kropinski, A. M., Alikhan, N.-F., Adriaenssens, E. M. & Consortium, I. R. ICTV virus taxonomy profile: Herelleviridae. The Journal of general virology 101, 362 (2020).

75. Nair, D. et al. Whole-genome sequencing of Staphylococcus aureus strain RN4220, a key laboratory strain used in virulence research, identifies mutations that affect not only virulence factors but also the fitness of the strain. Journal of bacteriology 193, 2332–2335 (2011).

76. Jones, M. J., Donegan, N. P., Mikheyeva, I. V. & Cheung, A. L. Improving transformation of Staphylococcus aureus belonging to the CC1, CC5 and CC8 clonal complexes. PLoS One 10, e0119487 (2015).

77. Zhang, D. et al. The use of a novel NanoLuc-based reporter phage for the detection of Escherichia coli O157: H7. Scientific reports 6, 33235 (2016).

78. Pulkkinen, E. M., Hinkley, T. C. & Nugen, S. R. Utilizing in vitro DNA assembly to engineer a synthetic T7 Nanoluc reporter phage for Escherichia coli detection. Integrative Biology 11, 63–68 (2019).

79. Jain, P. et al. Nanoluciferase reporter mycobacteriophage for sensitive and rapid detection of Mycobacterium tuberculosis drug susceptibility. Journal of Bacteriology 202, 10–1128 (2020).

80. Chen, Y. E., Fischbach, M. A. & Belkaid, Y. Skin microbiota–host interactions. Nature 553, 427–436 (2018).

81. Bryan, D., El-Shibiny, A., Hobbs, Z., Porter, J. & Kutter, E. M. Bacteriophage T4 Infection of Stationary Phase E. coli: Life after Log from a Phage Perspective. Frontiers in Microbiology 7. issn: 1664-302X. https://www.frontiersin.org/articles/10.3389/fmicb.2016.01391 (2016).

82. Ripp, S. & Miller, R. V. Dynamics of the pseudolysogenic response in slowly growing cells of Pseudomonas aeruginosa. Microbiology 144, 2225–2232. issn: 1465-2080. https://www.microbiologyresearch.org/content/journal/micro/10.1099/00221287-144-8-2225 (1998).

83. Maffei, E. et al. Phage Paride can kill dormant, antibiotic-tolerant cells of Pseudomonas aeruginosa by direct lytic replication. Nature Communications 15. issn: 2041-1723. 10.1038/s41467-023-44157-3 (Jan. 2024).

84. OpenAI, R. GPT-4 technical report. arXiv 2303.08774. View in Article (2023).

